# TRPM4 Couples Mechanical Force to Myogenic Constriction Throughout the Resistance Vasculature

**DOI:** 10.64898/2026.06.08.731006

**Authors:** Wanchun Zhu, Alfredo Sanchez–Solano, Boris Lavanderos, Shiyue Pan, Yumei Feng Earley, Scott Earley

## Abstract

**Background:** Myogenic tone is a fundamental property of resistance arteries that stabilizes tissue perfusion by coupling intraluminal pressure to smooth muscle cell (SMC) depolarization, Ca^2+^ influx, and vasoconstriction. TRPM4 (transient receptor potential melastatin 4) cation channels are required for this response in cerebral arteries, but whether TRPM4-dependent mechanotransduction is conserved across the broader resistance vasculature has remained unknown.

**Methods:** We combined droplet digital PCR, a newly generated *Trpm4*-Cre transgenic reporter mouse line, native-cell patch-clamp electrophysiology, pressure myography, selective pharmacological inhibition, and novel SMC-specific *Trpm4*-knockout (*Trpm4*-smKO) mice to define the expression, regulation, and functional importance of TRPM4 in cerebral, mesenteric, and skeletal muscle resistance arteries.

**Results:** *Trpm4* transcripts were detected in all three vascular beds, and genetic reporter-based mapping localized TRPM4 expression to SMCs in multiple organs. Using conventional whole-cell patch-clamp electrophysiology, we recorded cation currents activated by high intracellular [Ca^2+^] and sensitive to the selective TRPM4 blocker 4-chloro-2-(1-naphthyloxyacetamido) benzoic acid (NBA) in native SMCs from all three beds. In cells patch-clamped using the amphotericin B-perforated configuration, stretching the plasma membrane by applying negative pressure (-20 mmHg) through the patch pipette activated transient inward cation currents that were suppressed by NBA. The selective angiotensin II type 1 receptor (AT_1_R) blocker losartan also inhibited stretch-induced currents without affecting Ca^2+^-activated whole-cell TRPM4 currents, indicating that AT_1_R signaling is required for mechanotransduction in SMCs from all three vascular beds. In pressurized arteries with established myogenic tone, NBA produced reversible, concentration-dependent suppression of pressure-induced constriction of cerebral, mesenteric, and skeletal muscle arteries while sparing constriction induced by direct depolarization of SMCs with high (60 mM) extracellular [K^+^]. TRPM4-dependent whole-cell currents and stretch-induced cation currents were decreased in SMCs from *Trpm4*-smKO mice, and myogenic tone was essentially absent in all three vascular beds from these animals.

**Conclusions:** These findings show that TRPM4 is essential for pressure-induced SMC depolarization and myogenic constriction in the resistance vasculature.

## INTRODUCTION

Pressure-induced vasoconstriction, manifested as myogenic tone, is a defining feature of the small arteries and arterioles that regulate vascular resistance and tissue perfusion^1,2^. Myogenic tone arises from the integrated activity of ion channels that govern membrane potential and global intracellular Ca^2+^ concentration in vascular smooth muscle cells (SMCs)^3^. At physiological intraluminal pressures, mechanical stimuli engage signaling pathways in SMCs that evoke depolarizing currents, thereby increasing Ca^2+^ influx through voltage-dependent Ca_V_1.2 channels and activating the contractile apparatus to produce vasoconstriction^2,3^. Depolarization is counterbalanced by hyperpolarizing conductances mediated primarily by large-conductance Ca^2+^-activated K^+^ (BK) channels and voltage-dependent K^+^ (K_V_) channels, such that membrane potential and myogenic tone stabilize at steady-state levels^4–6^. Multiple ion channels have been implicated in the depolarizing arm of the myogenic response, including the transient receptor potential (TRP) channels TRPC6^7^, TRPV1^8,9^, TRPP1/PKD2^10,11^, and TRPM4^12,13^, as well as the Ca^2+^-activated Cl^−^ channel TMEM16A/ANO1^14^. Among these candidates, substantial prior evidence supports a critical role for TRPM4 in the development of myogenic tone in cerebral arteries^12,13,15,16^.

TRPM4 is a broadly expressed, Ca^2+^-activated, monovalent-cation–selective channel that is impermeable to Ca^2+^ ^17^. In SMCs from cerebral arteries, TRPM4 promotes membrane depolarization by conducting inward Na^+^ currents, thereby triggering Ca^2+^ influx and vasoconstriction^15,16^. Consistent with this role, prior studies have shown that TRPM4 expression and activity are required for pressure-induced SMC depolarization and myogenic tone in cerebral pial arteries^12,13,16^ and parenchymal arterioles^18,19^, and for autoregulation of cerebral blood flow *in vivo* in response to changes in perfusion pressure^20^. In patch-clamp experiments, stretching the plasma membranes of SMCs evokes transient inward cation currents (TICCs) attributable to TRPM4^13,15^. However, TRPM4 is not inherently mechanosensitive^13,21^. Instead, it is activated downstream of signaling pathways initiated by angiotensin II type 1 receptors (AT_1_Rs)^13,19,22^ and possibly other Gα_q/11_-coupled receptors^18^. In this model, receptor-driven phospholipase C (PLC) activity generates inositol 1,4,5-trisphosphate (IP_3_), which stimulates Ca^2+^ release from the sarcoplasmic reticulum (SR) through IP_3_ receptors (IP_3_Rs) to activate Ca^2+^-dependent TRPM4 channels at the plasma membrane^13^. Whether this TRPM4-dependent mechano-signaling mechanism is conserved across vascular beds has remained unclear.

Vascular beds differ in developmental origin, local environment, hemodynamic stress, and physiological function—differences that are accompanied by marked heterogeneity in the ion channels, receptors, and signaling pathways that govern membrane excitability and contractility^23^. Consequently, SMC phenotype is not uniform across the vasculature. Arteries supplying the brain, skeletal muscle, and the mesenteric circulation each operate under distinct regulatory constraints and could therefore employ different molecular mechanisms to sense intraluminal pressure and generate myogenic tone. Accordingly, although TRPM4-dependent signaling is a key determinant of myogenic reactivity in cerebral arteries, it remains unknown whether this mechanism is conserved in other resistance vessels. Resolving this issue is necessary to determine whether TRPM4 serves as a general regulator of arterial smooth muscle excitability or instead fulfills a specialized role in select vascular beds.

To test whether TRPM4-dependent signaling is a conserved mechanism governing myogenic tone across resistance vessels from distinct circulatory territories, we here compared the distribution and functional importance of TRPM4 in SMCs from cerebral, mesenteric, and skeletal muscle arteries using complementary pharmacological and genetic approaches, together with a newly developed reporter-based mapping approach. By integrating expression analyses with electrophysiological and pressure myography experiments, we sought to determine whether TRPM4 is a common downstream effector of AT_1_R-dependent mechanotransduction and serves as a broadly conserved regulator of arterial smooth muscle excitability and myogenic tone.

## RESULTS

### *Trpm4* is broadly expressed in arterial smooth muscle

To determine whether TRPM4 expression is conserved across resistance arteries from distinct circulations, we first quantified *Trpm4* transcript abundance in cerebral, mesenteric, and skeletal muscle arteries using droplet digital PCR (ddPCR). *Trpm4* mRNA was readily detected in all three vascular beds (Figure 1A). Transcript abundance was greater in mesenteric than in cerebral arteries, but did not differ significantly in the remaining pairwise comparisons.

**Figure 1:**
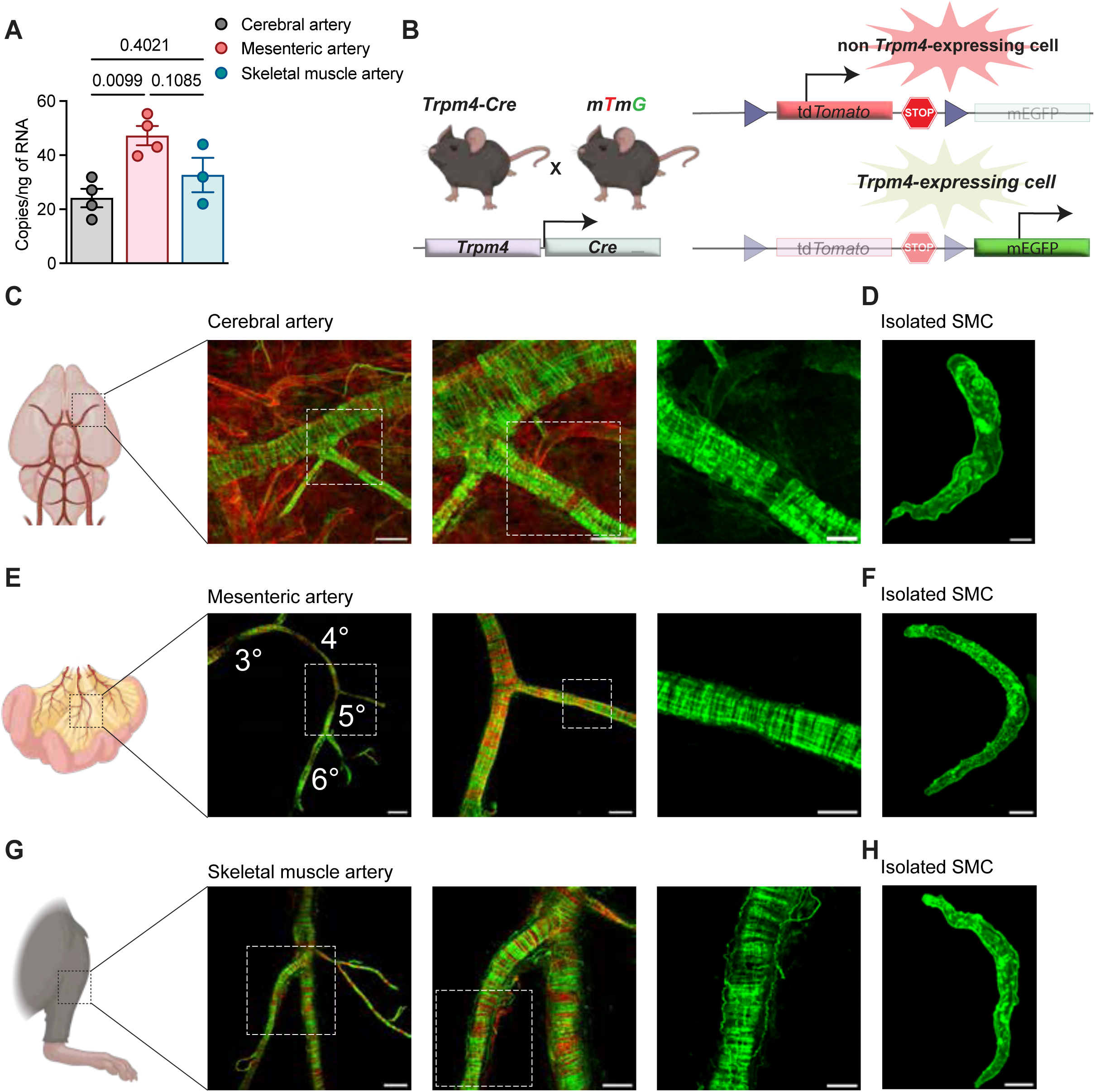
TRPM4 is expressed in arterial SMCs across vascular beds. **A,** ddPCR analysis of *Trpm4* transcript abundance (copies/ng RNA) in cerebral, mesenteric, and skeletal muscle arteries. Data are presented as means ± SEM (*N* = 3–4 animals; *P* = 0.0099 for cerebral vs. mesenteric arteries, *P* = 0.1085 for mesenteric vs. skeletal muscle arteries, and *P* = 0.4021 for cerebral vs. skeletal muscle arteries; one-way ANOVA with Tukey multiple-comparisons test). Symbols represent individual animals. **B**, Schematic illustrating the *Trpm4*-Cre reporter strategy. In *Trpm4*-Cre mice crossed with mT/mG reporter mice, cells without *Trpm4*-Cre activity express membrane-targeted tdTomato, whereas cells with *Trpm4*-Cre activity undergo Cre-mediated recombination, excision of the tdTomato cassette, and expression of membrane-targeted EGFP. **C**, Representative fluorescence images of cerebral arteries from *Trpm4*-Cre::mT/mG mice. Left, merged low-magnification view of the vascular network (tdTomato and mEGFP fluorescence). Middle, higher-magnification merged view of the boxed region shown on the left (tdTomato and mEGFP fluorescence). Right, zoomed-in mEGFP fluorescence image of the boxed region shown in the middle. Scale bars, 100 µm (left image), 50 µm (middle image), and 20 µm (right image). **D**, Representative enzymatically isolated cerebral artery SMC expressing mEGFP. Scale bar, 5 µm. **E**, Representative fluorescence images of mesenteric arteries from *Trpm4*-Cre::mT/mG mice. Left, merged low-magnification image. Middle, merged higher-magnification image. Right, mEGFP fluorescence. Scale bars, 500 µm (left), 200 µm (middle), and 100 µm (right). **F**, Representative enzymatically isolated mesenteric artery SMC expressing mEGFP. Scale bar, 10 µm. **G**, Representative fluorescence images of skeletal muscle arteries from *Trpm4*-Cre::mT/mG mice. Left, merged low-magnification image. Middle, merged higher-magnification image. Right, mEGFP fluorescence. Scale bars, 200 µm (left), 100 µm (middle), and 50 µm (right). **H**, Representative enzymatically isolated skeletal muscle artery SMC expressing mEGFP. Scale bar, 10 µm. All fluorescence images are representative of at least 3 independent vessel preparations, with consistent reporter expression observed across animals.

To localize TRPM4 expression at the cellular level, we developed a *Trpm4*-Cre knock-in transgenic mouse line in which Cre recombinase is expressed from the endogenous *Trpm4* locus. In this model, the native *Trpm4* stop codon in exon 25 was replaced with a P2A-Cre cassette using CRISPR/Cas-mediated genome engineering. Crossing these mice with the mTmG reporter line enabled identification of TRPM4-lineage cells by membrane-target GFP expression (Figure 1B). Robust EGFP fluorescence was observed throughout the medial layer of cerebral arteries in intact arteries from SMCs isolated from *Trpm4*-Cre::mTmG mice but was absent from the endothelial cell layer (Figure 1C and D). A similar pattern was observed for mesenteric (Figure 1E and F) and skeletal muscle (Figure 1G and H) arteries. These findings show that TRPM4 expression is not restricted to the cerebral circulation but is a shared feature of SMCs from multiple resistance artery beds.

### A common mechanotransduction signaling pathway regulates stretch-induced TRPM4 activity in SMCs across vascular beds

To determine whether TRPM4 channels are functional in peripheral vascular beds, we performed patch-clamp electrophysiology on native SMCs isolated from cerebral, mesenteric, and skeletal muscle arteries. Using the conventional whole-cell configuration, which allows free diffusion between the pipette solution and the intracellular compartment, we dialyzed cells with a high-Ca^2+^ pipette solution (200 μM free Ca^2+^) to directly activate TRPM4 channels. Under these conditions, voltage ramps (−100 to +100 mV from a holding potential of −60 mV) elicited outwardly rectifying cation currents that reversed near 0 mV. These currents developed within 20-30 s after rupture of the cell membrane under the patch pipette and reached a stable maximum amplitude by ∼100 s. Currents recorded from cerebral artery SMCs under these conditions were diminished by the selective TRPM4 inhibitor 4-chloro-2-(1-naphthyloxyacetamido) benzoic acid^24,25^ (NBA; 10 µM) (Figure 2A). To assess activation of TRPM4 by mechanical stimulation, we recorded TICCs^15^ using the perforated patch-clamp configuration, which allows us to control the membrane potential while preserving intracellular Ca^2+^ signaling^26^. Stretching the plasma membrane by application of negative pressure (−20 mmHg) through the patch pipette increased TICC activity, and both the frequency and amplitude of these mechanosensitive currents were significantly reduced by NBA (Figure 2B). NBA had similar inhibitory effects on Ca^2+^-activated whole-cell currents and stretch-induced TICCs in SMCs from mesenteric (Figure 2C and D) and skeletal muscle (Figure 2E and F) arteries.

**Figure 2.**
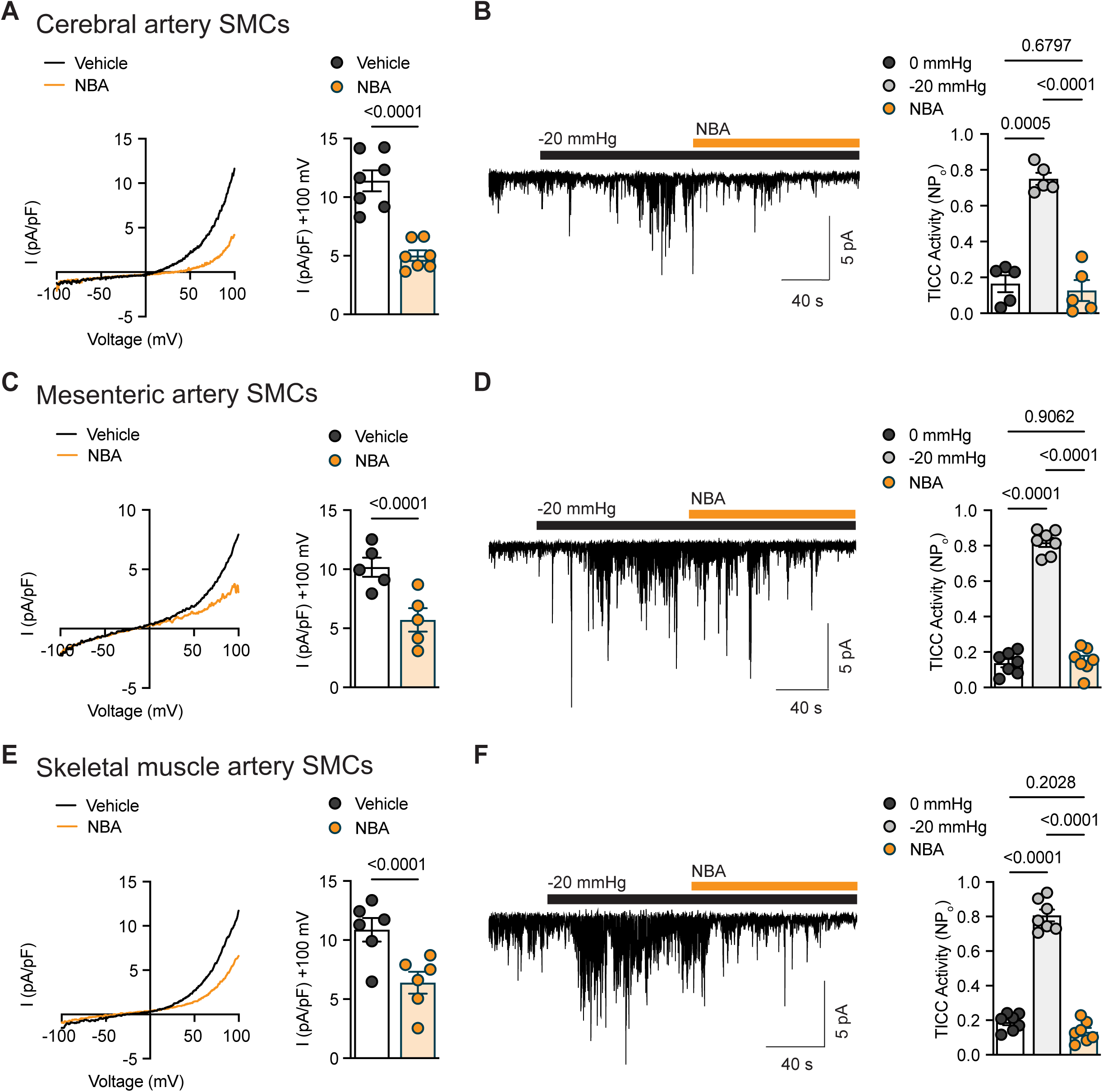
Stretch-activated, NBA-sensitive TRPM4 currents are present in SMCs across vascular beds. **A**, Conventional whole-cell patch-clamp recordings from cerebral artery SMCs. Left, representative current–voltage (I–V) relationship under vehicle conditions and after application of NBA (10 µM). Right, summary current density measured at +100 mV. Data are means ± SEM (n = 7 cells from 6 mice; *P* < 0.0001; paired *t*-test). Symbols represent individual cells. **B**, Perforated-patch recordings from cerebral artery SMCs. Left, representative recording showing TICC activity at 0 mmHg, during membrane stretch induced by −20 mmHg suction, and after NBA. Right, summary TICC activity expressed as NP_o_. Data are means ± SEM (n = 5 cells from 4 mice; *P* = 0.0005 for 0 mmHg vs. −20 mmHg, *P* < 0.0001 for −20 mmHg vs. NBA, and *P* = 0.6797 for 0 mmHg vs. NBA; repeated-measures one-way ANOVA with Tukey test). Symbols represent individual cells. **C**, Conventional whole-cell recordings from mesenteric artery SMCs. Left, representative I–V relationship under vehicle conditions and after NBA. Right, summary current density at +100 mV. Data are means ± SEM (n = 5 cells from 5 mice; *P* < 0.0001; paired *t*-test). **D**, Perforated-patch recordings from mesenteric artery SMCs. Left, representative recording showing stretch-induced TICC activity at 0 mmHg, −20 mmHg, and after NBA. Right, summary TICC activity. Data are means ± SEM (n = 7 cells from 6 mice; *P* < 0.0001 for 0 mmHg versus −20 mmHg, *P* < 0.0001 for −20 mmHg vs. NBA, and *P* = 0.9062 for 0 mmHg vs. NBA; repeated-measures one-way ANOVA with Tukey test). **E**, Conventional whole-cell recordings from skeletal muscle artery SMCs. Left, representative I–V relationship under vehicle conditions and after NBA. Right, summary current density at +100 mV. Data are means ± SEM (n = 6 cells from 4 mice; *P* < 0.0001; paired *t*-test). **F**, Perforated-patch recordings from skeletal muscle artery SMCs. Left, representative recording showing stretch-induced TICC activity at 0 mmHg, −20 mmHg, and after NBA. Right, summary TICC activity. Data are means ± SEM (n = 7 cells from 3 mice; *P* < 0.0001 for 0 mmHg vs. −20 mmHg, *P* < 0.0001 for −20 mmHg vs. NBA, and *P* = 0.2028 for 0 mmHg vs. NBA; repeated-measures one-way ANOVA with Tukey test).

Stretching the plasma membrane triggers an AT_1_R-initiated, PLC-dependent signaling cascade that activates TRPM4 channels in SMCs from cerebral arteries and arterioles^13,19,22^. We used the selective AT_1_R antagonist losartan (10 µM) to determine whether the same is true in other vascular beds. Whole-cell patch-clamp recordings showed that losartan did not affect Ca^2+^-activated whole-cell TRPM4 currents in SMCs from cerebral arteries (Supplemental Figure S1A), indicating that it does not directly inhibit TRPM4 channels. In contrast, perforated patch-clamp recordings revealed that losartan markedly suppressed negative pressure (−20 mmHg)-induced currents in these cells (Supplemental Figure S1B). Losartan had similar effects in SMCs from mesenteric (Supplemental Figure S1C and D) and skeletal muscle (Supplemental Figure S1E and F) arteries, indicating that AT_1_R signaling is required for stretch-induced TRPM4 activation in these peripheral vascular beds. Together, these results demonstrate that TRPM4 channels are functionally active across multiple vascular beds and are regulated by a conserved AT_1_R-dependent mechanosensing signaling pathway.

### Selective pharmacological inhibition of TRPM4 reversibly eliminates myogenic tone

To determine the effects of NBA on myogenic tone, we pressurized cannulated cerebral arteries to physiological levels (∼40-80 mmHg) using standard pressure myography methods^27^ and allowed them to develop spontaneous myogenic tone. Vessels were then superfused with a single concentration of NBA (range, 10 nM to 30 µM) in a noncumulative design, with continuous recording of luminal diameter. At the end of each experiment, passive diameter was determined by superfusing with a Ca^2+^-free solution, and myogenic tone was expressed relative to passive diameter. Higher concentrations of NBA produced rapid and reversible vasodilation, with robust effects evident at 10 µM (Figure 3A). NBA caused modest dilation at 1 µM and near-maximal dilation at 30 µM, yielding an IC_50_ (50% inhibitory concentration) of 2.6 µM in cerebral arteries (Figure 3B). This value is comparable to IC_50_ values obtained using fluorescence-based *in vitro* screening assays (IC_50_ for transfected HEK293 cells ≈ 1.3 μM; IC_50_ for HCT116 human colorectal cancer cells ≈ 2.7 μM)^24,25^. Similar concentration-dependent inhibition of myogenic tone by NBA was observed in mesenteric (Figure 3C and D) and skeletal muscle (Figure 3E and F) arteries, with IC_50_ values of 6.4 µM and 8.9 µM, respectively.

**Figure 3.**
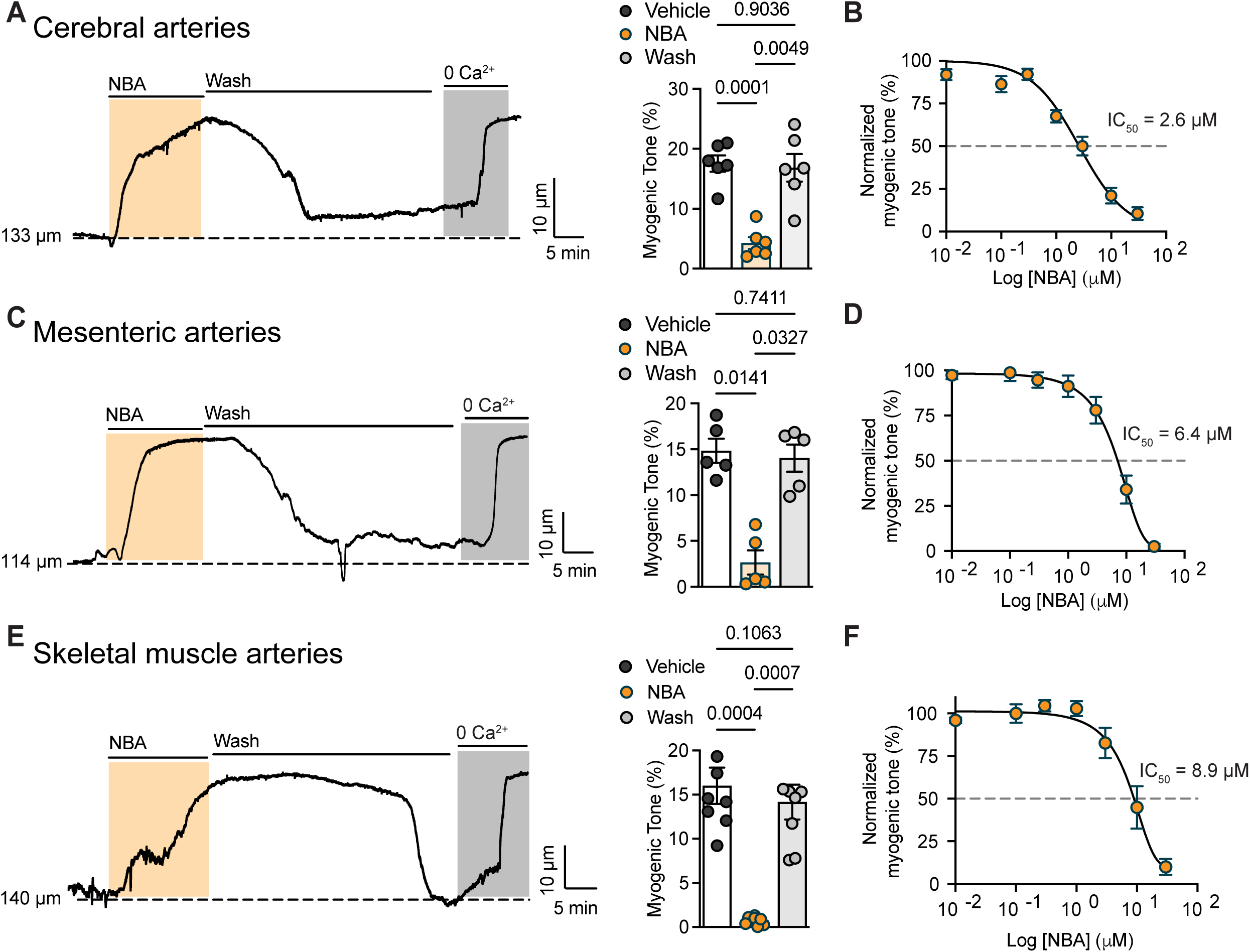
Selective pharmacological inhibition of TRPM4 reversibly eliminates myogenic tone. **A,** Representative diameter trace from a pressurized cerebral artery showing inhibition of myogenic tone by NBA (10 µM), recovery after washout, and maximal dilation in Ca^2+^-free solution. Dashed line indicates initial diameter. Right, summary data for myogenic tone under vehicle conditions, after NBA, and after washout. Data are means ± SEM (n = 6 arteries from 5 mice; *P* = 0.0001 for vehicle vs. NBA, *P* = 0.0049 for NBA vs. wash, and *P* = 0.9036 for vehicle vs. wash; repeated-measures one-way ANOVA with Tukey test). Symbols represent individual arteries. **B**, Concentration–response curve for inhibition of myogenic tone by NBA in isolated cerebral arteries. Myogenic tone is normalized to tone in the absence of NBA. Data are means ± SEM (n = 37 arteries from 22 mice). Curve was fit by nonlinear regression; IC_50_ = 2.6 µM. **C**, Representative diameter trace from a pressurized mesenteric artery showing inhibition of myogenic tone by NBA (10 µM), recovery after washout, and maximal dilation in Ca^2+^-free solution. Dashed line indicates initial diameter. Right, summary data for myogenic tone under vehicle conditions, after NBA, and after washout. Data are means ± SEM (n = 5 arteries from 4 mice; *P* = 0.0141 for vehicle vs. NBA, *P* = 0.0327 for NBA vs. wash, and *P* = 0.7411 for vehicle vs. wash by repeated-measures one-way ANOVA with Tukey test). Symbols represent individual arteries. **D**, Concentration–response curve for inhibition of myogenic tone by NBA in isolated mesenteric arteries. Myogenic tone is normalized to tone in the absence of NBA. Data are means ± SEM (n = 5-8 arteries from 7 mice). Curve was fit by nonlinear regression; IC_50_ = 6.4 µM. **E**, Representative diameter trace from a pressurized skeletal muscle artery showing inhibition of myogenic tone by NBA (10 µM), recovery after washout, and maximal dilation in Ca^2+^-free solution. Dashed line indicates initial diameter. Right, summary data for myogenic tone under vehicle conditions, after NBA, and after washout. Data are means ± SEM (n = 8 arteries from 8 mice; *P* = 0.0004 for vehicle vs. NBA, *P* = 0.0007 for NBA vs. wash, and *P* = 0.1063 for vehicle vs. wash; repeated-measures one-way ANOVA with Tukey test). Symbols represent individual arteries. **F**, Concentration–response curve for inhibition of myogenic tone by NBA in isolated skeletal muscle arteries. Myogenic tone is normalized to tone in the absence of NBA. Data are means ± SEM (n = 6 arteries from 4 mice). Curve was fit by nonlinear regression; IC_50_ = 8.9 µM.

We next examined the effect of NBA on the development of myogenic tone over a broad pressure range. Cannulated arteries were subjected to stepwise increases in intraluminal pressure (5-140 mmHg), and active diameter was measured at each pressure. Pressure-diameter relationships were then reexamined in the presence of NBA (10 µM) and, finally, in Ca^2+^-free solution to establish the passive diameter at each pressure. Myogenic tone was calculated at each pressure as the difference between passive and active diameters, normalized to the passive diameter. Under control conditions, cerebral arteries developed robust pressure-induced constriction beginning at approximately 40 mmHg (Figure 4A). In contrast, NBA nearly abolished constriction across the pressure range, markedly reducing myogenic tone relative to vehicle treatment (Figure 4A). Importantly, constriction evoked by superfusion with 60 mM K^+^ was unaffected by NBA (Figure 4B), indicating that the drug does not inhibit voltage-dependent Ca^2+^ entry or directly disrupt the contractile apparatus. NBA also strongly suppressed the development of myogenic tone in mesenteric (Figure 4C and D) and skeletal muscle (Figure 4E and F) arteries. Together, these findings demonstrate that selective pharmacological blockade of TRPM4 activity inhibits the development of myogenic tone in cerebral, mesenteric, and skeletal muscle resistance arteries.

**Figure 4.**
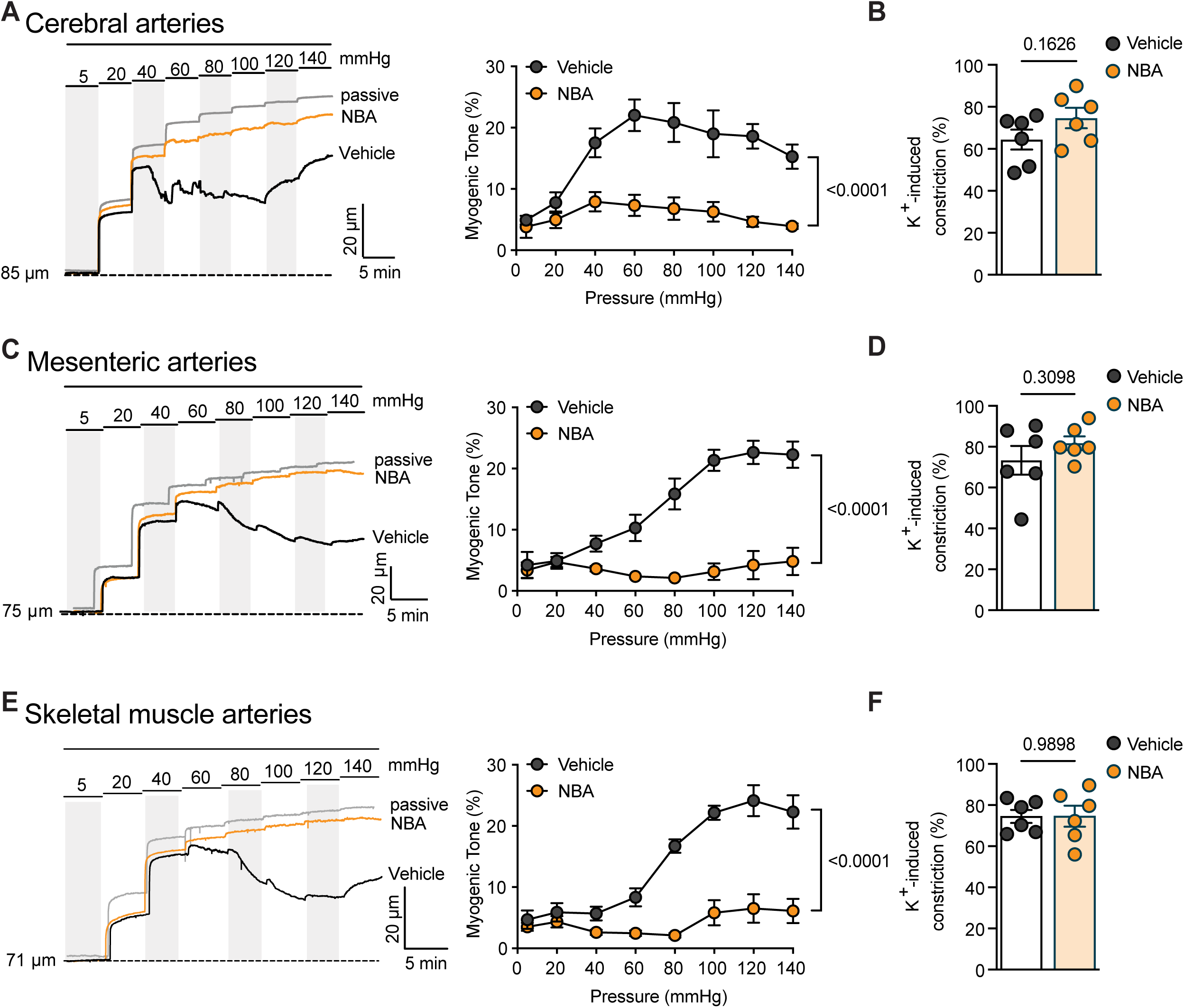
Selective TRPM4 blockade prevents the development of myogenic tone without impairing direct depolarization-induced vasoconstriction. **A**, Representative luminal diameter recordings from isolated cerebral arteries exposed to stepwise increases in intraluminal pressure under vehicle conditions, in the presence of NBA (10 µM), and after removal of extracellular Ca^2+^ (to determine passive diameter). Right, summary pressure–myogenic tone relationship. Data are means ± SEM (n = 6 arteries from 6 mice; *P* < 0.0001 for vehicle vs. NBA; two-way ANOVA). **B**, Summary data showing constriction of cerebral arteries in response to 60 mM K^+^ under vehicle conditions and in the presence of NBA. Data are means ± SEM (n = 6 arteries from 6 mice; *P* = 0.1626; paired *t*-test). Symbols represent individual arteries. **C**, Representative luminal diameter recordings from isolated mesenteric arteries exposed to stepwise increases in intraluminal pressure under vehicle conditions, in the presence of NBA, and in Ca^2+^-free solution (to determine passive diameter). Right, summary pressure–myogenic tone relationship. Data are means ± SEM (n = 6 arteries from 5 mice; *P* < 0.0001 for vehicle vs. NBA; two-way ANOVA). **D**, Summary data showing 60 mM K^+^-induced constriction of mesenteric arteries under vehicle conditions and in the presence of NBA. Data are means ± SEM (n = 6 arteries from 5 mice; *P* = 0.3098; paired *t*-test). Symbols represent individual arteries. **E**, Representative luminal diameter recordings from isolated skeletal muscle arteries exposed to stepwise increases in intraluminal pressure under vehicle conditions, in the presence of NBA, and in Ca^2+^-free solution (to determine passive diameter). Right, summary pressure–myogenic tone relationship. Data are means ± SEM (n = 6 arteries from 4 mice; *P* < 0.0001 for vehicle vs. NBA; two-way ANOVA). **F**, Summary data showing 60 mM K^+^-induced constriction of skeletal muscle arteries under vehicle conditions and in the presence of NBA. Data are means ± SEM (n = 6 arteries from 4 mice; *P* = 0.9898 by paired *t*-test). Symbols represent individual arteries.

### Smooth muscle *Trpm4* deletion suppresses Ca^2+^- and stretch-activated TRPM4 currents

To complement our pharmacological studies, we generated smooth muscle *Trpm4*-knockout (*Trpm4*-smKO) mice by crossing newly generated *Trpm4^fl/fl^* mice, in which exon 10 is flanked by loxP sites, with SM22-Cre animals (Figure 5A). *Trpm4*-smKO mice were viable, normal in appearance and growth rate, and exhibited the expected Mendelian frequency. Analysis of transcript levels using ddPCR revealed a marked reduction in *Trpm4* expression in cerebral, mesenteric, and skeletal muscle arteries from *Trpm4*-smKO mice compared with *Trpm4*^fl/fl^ controls (Figure 5B). In contrast, transcript levels of *Trpv1*, *Trpc6*, *Ano1,* and *Pkd2* were unchanged (Figure 5C), supporting the specificity of *Trpm4* deletion and the absence of compensatory changes in the expression of related channels.

**Figure 5.**
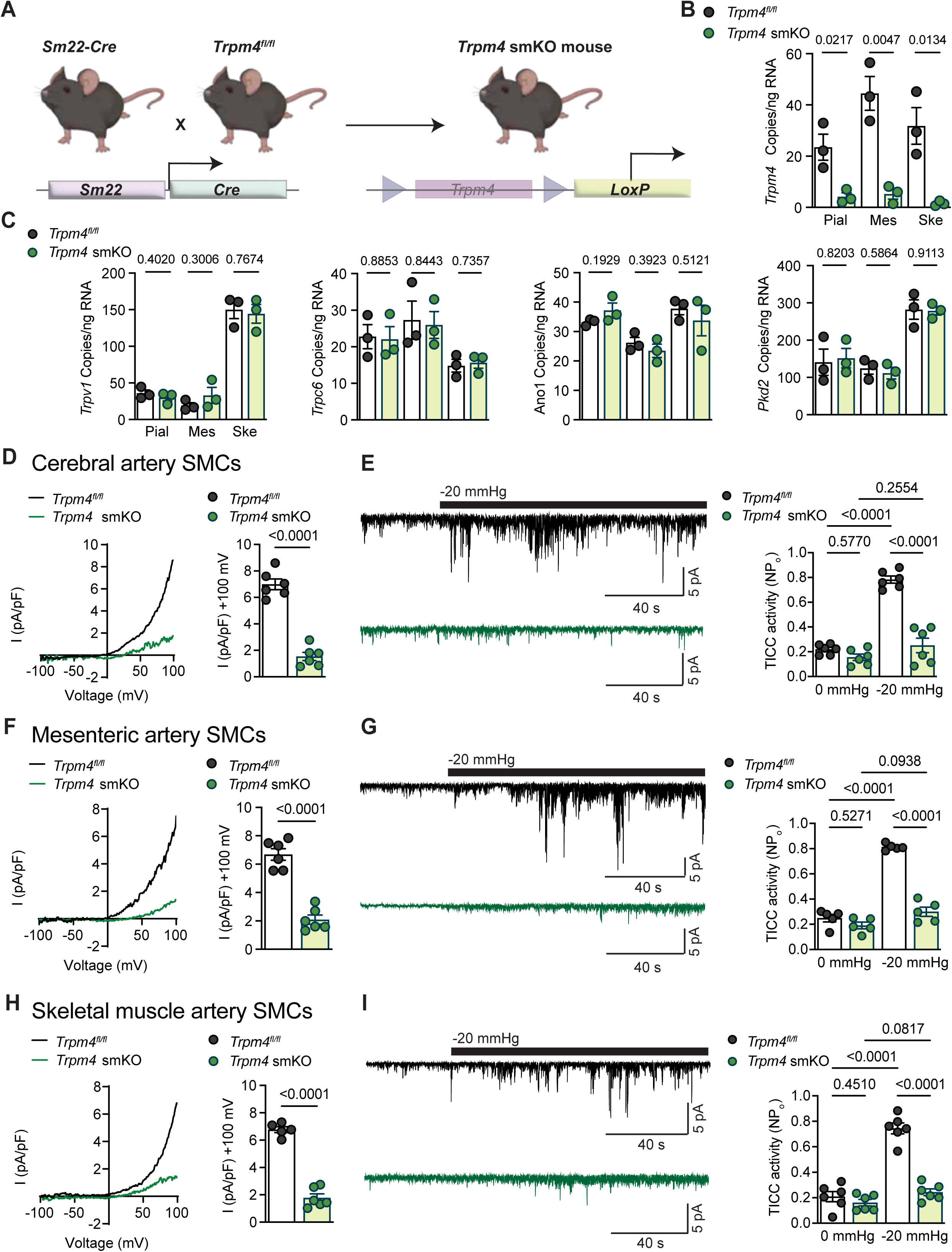
Smooth muscle-specific *Trpm4* deletion suppresses Ca^2+^-activated TRPM4 currents and stretch-induced TICC activity. **A,** Schematic illustrating the breeding strategy used to generate smooth muscle-specific *Trpm4*-KO (*Trpm4-*smKO) mice by crossing Sm22-Cre mice with *Trpm4^fl/fl^* mice. **B**, ddPCR analysis showing reduced *Trpm4* transcript abundance (copies/ng RNA) in cerebral, mesenteric, and skeletal muscle arteries from *Trpm4*-smKO mice compared with *Trpm4^fl/fl^* controls. Data are presented as means ± SEM (N = 3 animals per group; *P* = 0.0217 for cerebral arteries, *P* = 0.0047 for mesenteric arteries, and *P* = 0.0134 for skeletal muscle arteries from *Trpm4^fl/fl^*vs.*Trpm4*-smKO mice; unpaired *t*-test). Symbols represent individual animals. **C**, ddPCR analysis of *Trpv1*, *Trpc6*, *Ano1*, and *Pkd2* transcript abundance (copies/ng RNA) in cerebral, mesenteric, and skeletal muscle arteries from *Trpm4^fl/fl^* and *Trpm4*-smKO mice. Data are presented as means ± SEM (N = 3 animals per group). *P*-values for the indicated comparisons were determined by unpaired *t*-tests. Symbols represent individual animals. **D**, Conventional whole-cell patch-clamp recordings from cerebral artery SMCs isolated from *Trpm4^fl/fl^* and *Trpm4*-smKO mice. Left, representative current–voltage relationships. Right, summary current density measured at +100 mV. Data are means ± SEM (n = 6 cells from 4 mice per group; *P* < 0.0001; unpaired *t*-test). Symbols represent individual cells. **E**, Perforated-patch recordings from cerebral artery SMCs isolated from *Trpm4^fl/fl^* and *Trpm4*-smKO mice. Left, representative traces showing TICC activity during membrane stretch induced by −20 mmHg suction. Right, summary TICC activity expressed as NP_o_. Data are means ± SEM (n = 6 cells from 3–4 mice per group). *P*-values for indicated comparisons were determined by two-way ANOVA with Tukey test. Symbols represent individual cells. **F**, Conventional whole-cell patch-clamp recordings from mesenteric artery SMCs isolated from *Trpm4^fl/fl^*and *Trpm4*-smKO mice. Left, representative current–voltage (I–V) relationships. Right, summary current density measured at +100 mV. Data are means ± SEM (n = 6 cells from 2–3 mice per group; *P* < 0.0001; unpaired *t*-test). Symbols represent individual cells. **G**, Perforated-patch recordings from mesenteric artery SMCs isolated from *Trpm4^fl/fl^* and *Trpm4*-smKO mice. Left, representative traces showing stretch-induced TICC activity during −20 mmHg suction. Right, summary of TICC activity expressed as NP_o_. Data are means ± SEM (n = 5 cells from 2–3 mice per group). *P*-values for indicated comparisons were determined by two-way ANOVA Tukey test. Symbols represent individual cells. **H**, Conventional whole-cell patch-clamp recordings from skeletal muscle artery SMCs isolated from *Trpm4^fl/fl^* and *Trpm4*-smKO mice. Left, representative I–V relationships. Right, summary current density measured at +100 mV. Data are means ± SEM (n = 5–6 cells from 3 mice per group; *P* < 0.0001; unpaired *t*-test). Symbols represent individual cells. **I**, Perforated-patch recordings from skeletal muscle artery SMCs isolated from *Trpm4^fl/fl^*and *Trpm4*-smKO mice. Left, representative traces showing stretch-induced TICC activity during −20 mmHg suction. Right, summary TICC activity expressed as NP_o_. Data are means ± SEM (n = 6 cells from 4 mice per group). *P*-values for indicated comparisons were determined by two-way ANOVA with Tukey test. Symbols represent individual cells.

We next used patch-clamp electrophysiology to assess the functional consequences of smooth muscle-specific *Trpm4* deletion. In conventional whole-cell recordings with a high-Ca^2+^ pipette solution, NBA-sensitive TRPM4 currents were significantly reduced in cerebral artery SMCs from *Trpm4*-smKO mice compared with *Trpm4^fl/fl^* controls (Figure 5D). In perforated patch-clamp recordings, stretch-induced TICC activity evoked by negative pressure (−20 mmHg) was also diminished in cerebral artery SMCs from *Trpm4*-smKO mice (Figure 5E). Similar reductions in Ca^2+^-activated and stretch-induced cation currents were obtained using SMCs from mesenteric (Figure 5F and G) and skeletal muscle (Figure 5H and I) arteries. These findings demonstrate that smooth muscle deletion of *Trpm4* effectively suppresses Ca^2+^- and stretch-induced TRPM4 channel activity across multiple vascular beds.

### Smooth muscle deletion of *Trpm4* suppresses myogenic tone across vascular beds

To determine whether TRPM4 expressed in SMCs is required for myogenic reactivity, we compared pressure-induced constriction in arteries isolated from *Trpm4^fl/fl^* and *Trpm4*-smKO mice. Stepwise increases in intraluminal pressure elicited robust myogenic constriction in cerebral arteries from *Trpm4^fl/fl^* mice, whereas they caused barely detectable constriction in cerebral arteries from *Trpm4-*smKO mice (Figure 6A), which exhibited significantly lower myogenic tone than controls (Figure 6B). In contrast, constriction evoked by directly depolarizing SMCs with high extracellular K^+^ did not differ between groups (Figure 6C), indicating that loss of *Trpm4* expression does not impair the fundamental contractile machinery or voltage-dependent Ca^2+^ entry. Similar findings were obtained using mesenteric (Figure 6D–F) and skeletal muscle (Figure 6G–I) arteries. These findings support the conclusion that TRPM4 is required for the development of myogenic tone across vascular beds.

**Figure 6.**
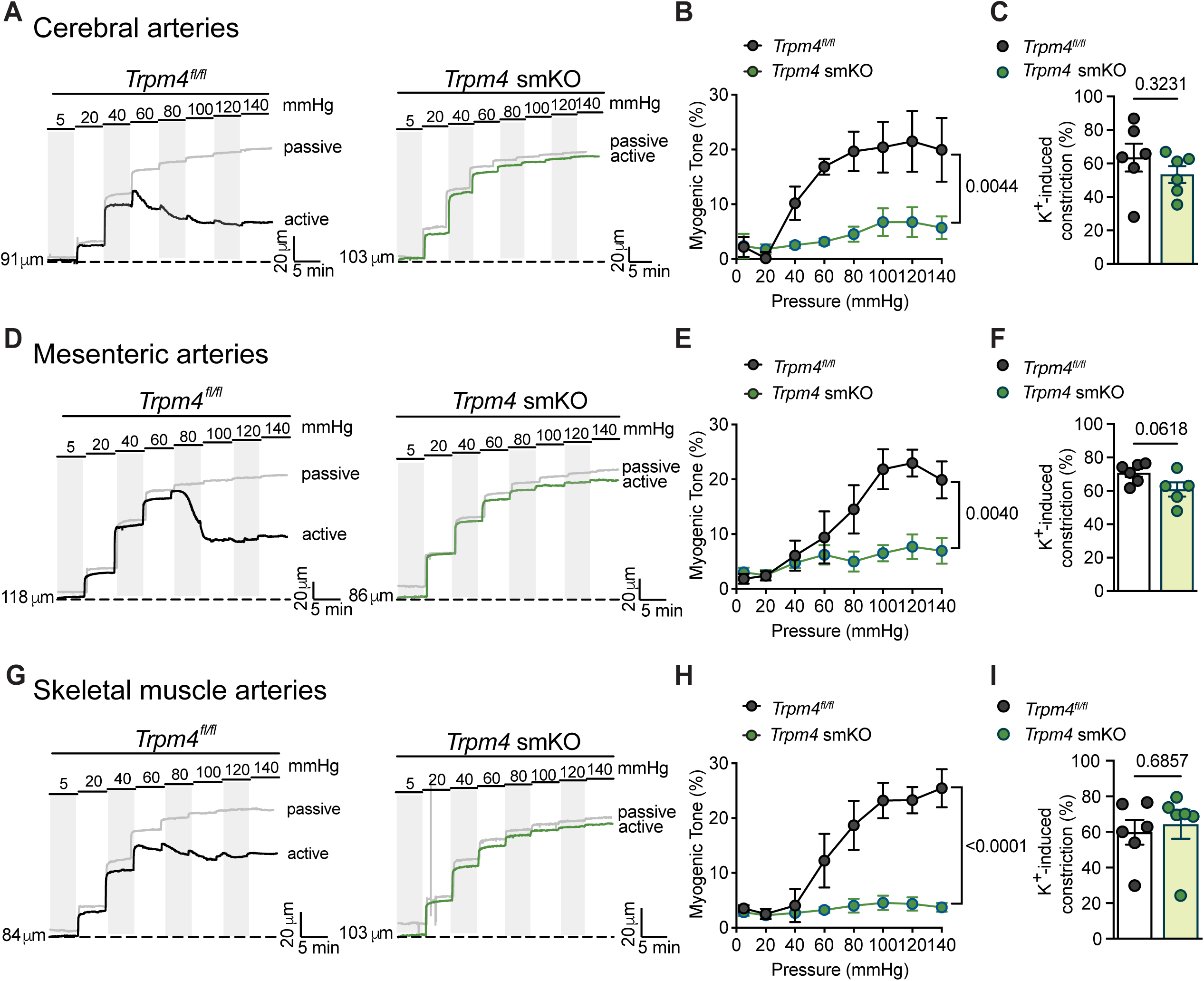
Smooth muscle-specific *Trpm4* deletion attenuates myogenic tone across vascular beds without impairing direct depolarization-induced constriction. A, Representative luminal diameter recordings from isolated cerebral arteries from *Trpm4^fl/fl^*and *Trpm4*-smKO mice exposed to stepwise increases in intraluminal pressure in Ca^2+^-containing (active) and Ca^2+^-free (passive) solution. **B**, Summary pressure–myogenic tone relationship for cerebral arteries. Data are means ± SEM (n = 6 arteries from 5 mice per group; *P* = 0.0044 for *Trpm4^fl/fl^* vs. *Trpm4*-smKO; repeated measures two-way ANOVA). **C**, Summary data showing constriction of cerebral arteries from *Trpm4^fl/fl^* and *Trpm4*-smKO mice in response to elevated extracellular K^+^ (60 mM). Data are means ± SEM (n = 6 arteries from 5 mice per group; *P* = 0.3231; unpaired *t*-test). Symbols represent individual arteries. **D**, Representative luminal diameter recordings of mesenteric arteries isolated from *Trpm4^fl/fl^* and *Trpm4*-smKO mice exposed to stepwise increases in intraluminal pressure under active and passive conditions. **E**, Summary pressure–myogenic tone relationship for mesenteric arteries. Data are means ± SEM (n = 5–6 arteries from 3–5 mice per group; *P* = 0.0040 for *Trpm4^fl/fl^* vs. *Trpm4*-smKO; repeated measures two-way ANOVA). **F**, Summary data showing constriction of mesenteric arteries from Trpm*4^fl/fl^* and *Trpm4*-smKO mice in response to elevated extracellular K^+^ (60 mM). Data are means ± SEM (n = 5–6 arteries from 3–5 mice per group; *P* = 0.0618; unpaired *t*-test). Symbols represent individual arteries. **G**, Representative luminal diameter recordings from skeletal muscle arteries isolated from *Trpm4^fl/fl^* and *Trpm4*-smKO mice exposed to stepwise increases in intraluminal pressure under active and passive conditions. **H**, Summary pressure–myogenic tone relationship for skeletal muscle arteries. Data are means ± SEM (n = 6 arteries from 4–6 mice per group; *P* < 0.0001 for *Trpm4^fl/fl^*vs. *Trpm4*-smKO mice; repeated measures two-way ANOVA). **I**, Summary data showing constriction of skeletal muscle arteries from *Trpm4^fl/fl^*and *Trpm4*-smKO mice in response to elevated extracellular K^+^ (60 mM). Data are means ± SEM (n = 6 arteries from 4–6 mice per group; *P* = 0.6857; unpaired *t*-test). Symbols represent individual arteries.

## DISCUSSION

The present study demonstrates a broad requirement for TRPM4 in the development of arterial myogenic tone throughout the vasculature. Although previous work established that TRPM4 is essential for pressure-induced constriction of cerebral arteries and arterioles, whether this mechanism extended beyond the brain remained unknown. Using a newly developed *Trpm4* reporter mouse, we show that TRPM4 is expressed in SMCs across multiple vascular beds. Electrophysiological studies focusing on SMCs from cerebral, mesenteric, and skeletal muscle arteries demonstrate that TRPM4 mediates stretch-activated inward cation currents in each vascular territory. Further, selective pharmacological inhibition of TRPM4 universally abolished myogenic tone, an effect that was replicated by smooth muscle deletion of *Trpm4*. These findings identify TRPM4 as a common and essential component of the myogenic apparatus across the vasculature.

Pressure-induced vasoconstriction arises from SMC depolarization and subsequent Ca^2+^ entry. Multiple ion channels have been proposed to initiate or amplify this depolarizing signal, but their contributions appear to vary substantially among vascular beds. For example, antisense-mediated knockdown of TRPC6 attenuates pressure-induced constriction of cerebral arteries^7^, whereas global *Trpc6* deletion does not impair the development of myogenic tone in mesenteric arteries^28^. This difference may reflect regional specialization of myogenic signaling or compensatory upregulation of *Trpc3* in *Trpc6-*KO mice^29,30^. Similarly, TRPV1 channels are highly expressed in skeletal muscle arterioles, where they have been proposed to contribute to myogenic tone in parallel with TRPM4^8,31^, whereas smooth muscle-specific deletion of *Pkd2* (encoding the TRP channel TRPP1) blunts myogenic tone in skeletal muscle arteries but not in mesenteric arteries^11^. Against this background of regional heterogeneity, the present study identifies TRPM4 as a broadly conserved determinant of myogenic tone. Using both selective pharmacological inhibition and smooth muscle genetic deletion, we show that TRPM4 channel activity is required for the full development of myogenic tone in all vascular beds tested.

These findings do not exclude contributions from other ion channels. Rather, they suggest that TRPM4 serves as a common depolarizing effector on which diverse, vascular bed-specific upstream mechanisms converge. In cerebral arteries, TRPC6 has been proposed to operate in series with TRPM4^13^, consistent with a model in which stretch-dependent SR Ca^2+^ release activates TRPM4 to amplify membrane depolarization. In skeletal muscle arterioles, TRPM4 channel blockade revealed TRPV1-dependent effects^9^, suggesting that TRPM4 and TRPV1 can act in parallel within the same vascular segment. Given the central role of TRPM4 as a common depolarizing effector elucidated here, future investigations should focus on how other channels implicated in the myogenic response influence TRPM4 expression, activity, and membrane localization.

Ion channels in the plasma membrane can be directly gated by mechanical stimuli or indirectly activated by signaling pathways initiated by force-sensitive receptors. Current evidence does not support the inherent mechanosensitivity of TRPM4 or of any other channel implicated in pressure-induced SMC depolarization, including TRPC6^7^, TRPV1^8^, TRPP1/PKD2^10,11^, and TMEM16A/ANO1^14^. Indeed, a systematic analysis of 11 TRP channels from 6 subfamilies, including TRPM4, TRPC6, TRPV1, and TRPP2/PKD2L1, concluded that none were intrinsically mechanosensitive^21^. Several reports provide compelling evidence that AT_1_Rs, and possibly other G-protein-coupled receptors that signal through Gα_q/11_ subunits, are inherently mechanosensitive and function as sensors of membrane stretch in SMCs^28,32,33^. PLC-dependent signaling pathways downstream of AT_1_R generate IP_3_ and diacylglycerol (DAG). IP_3_ stimulates Ca^2+^ release from IP_3_Rs on the SR to activate Ca^2+^-sensitive TRPM4 channels^13^, and DAG stimulates protein kinase C-dependent trafficking of TRPM4 to the plasma membrane^34,35^. The present data suggest that this signaling architecture is conserved across cerebral, mesenteric, and skeletal muscle arteries, indicating a common mechanosensory mechanism. Prior work also provides evidence that TRPC6 channels may be activated by the same PLC-driven pathway through DAG^32,36^, although this has not been definitively established in contractile SMCs. The upstream mechanisms that activate TRPV1, TRPP1/PKD2, and TMEM16A/ANO1 in response to membrane stretch are not known. These mechanotransduction pathways and their connections with AT_1_R and TRPM4 represent important targets for future investigation.

Losartan and other angiotensin receptor blocking drugs are widely used as antihypertensive agents. Our finding that losartan suppressed stretch-induced, AT_1_R-dependent activation of TRPM4 in arterial SMCs in multiple organs raises the possibility that inhibition of myogenic tone contributes to the reduction in peripheral vascular resistance produced by these drugs. Although the blood pressure-lowering actions of angiotensin receptor blockers are multifactorial, reduced AT_1_R–TRPM4 coupling in resistance arteries could represent an additional mechanism through which these agents blunt pressure-dependent vasoconstriction. The current findings may also have implications for small vessel diseases involving altered extracellular matrix–smooth muscle mechanotransduction. Prior studies indicate that cerebral vascular dysfunction in mouse models of Gould syndrome^37^, a rare autosomal dominant multisystem disorder caused by mutations in *COL4A1 or COL4A2*, encoding the extracellular matrix protein collagen IV, results in part from impaired smooth muscle TRPM4 activity^38,39^. Future studies should determine whether similar defects in TRPM4-dependent myogenic signaling occur in peripheral resistance arteries, which also contain collagen IV as a major component of the vascular extracellular matrix.

In summary, this study identifies smooth muscle TRPM4 channels as conserved effectors of pressure-induced vasoconstriction across multiple resistance arteries. Although smooth muscle cells from different vascular beds exhibit substantial heterogeneity in signaling architecture and contractile regulation, our findings indicate that TRPM4 provides a common depolarizing mechanism that is required for the full development of myogenic tone.

## MATERIALS AND METHODS

### Chemicals and Reagents

Unless otherwise specified, chemicals and other reagents were obtained from Sigma-Aldrich, Inc (St. Louis, MO, USA).

### Animals

All animal studies were performed in accordance with the guidelines of the University Committee on Animal Resources (UCAR) of the University of Rochester. All mice were maintained in individually ventilated cages (≤5 mice/cage) and housed in 12 hr/12 hr day/night cycle with *ad libitum* access to food (standard chow) and water. Male and female mice aged 3-4 mo were used for all experiments. Unless otherwise specified, all mice were maintained on a C57BL/6J background and purchased from Jackson Laboratories (Bar Harbor, ME, USA).

### Generation of *Trpm4*-mT/mG reporter mice

To identify *Trpm4*-expressing cells, we employed a *Trpm4*-P2A-Cre knock-in mouse line, generated in a C57BL/6NTac background by replacing the endogenous *Trpm4* stop codon in exon 25 with a P2A-Cre cassette using CRISPR/Cas-mediated genome engineering. The *Trpm4*-P2A-Cre line was crossed with the *Rosa26-*mTmG reporter strain such that cells without Cre-mediated recombination expressed membrane-targeted tdTomato, whereas cells expressing *Trpm4*-driven Cre underwent recombination and expressed membrane-targeted GFP, enabling visualization of TRPM4-expressing cells in intact tissues and isolated vessels.

### Generation of *Trpm4*-smKO mice

*Trpm4^fl/fl^* mice were generated by flanking exon 10 with loxP sites. Exon 10 was selected as the conditional knockout region because its deletion is predicted to cause loss of function through a frameshift, producing a truncated transcript with a premature stop codon that is expected to undergo nonsense-mediated decay. The effective conditional knockout region is approximately 1113 bp in length and does not overlap with any other known genes. *Trpm4-*smKO mice were generated by crossing *Trpm4^fl/fl^* mice with transgenic mice expressing Cre recombinase under the control of the smooth muscle-specific SM22α (*Tagln*) promoter (SM22-Cre).

### Isolation of arteries and SMCs

Mice were anesthetized with isoflurane (Baxter Healthcare) and then euthanized by decapitation and exsanguination. Brains, mesenteries, and hind limbs were isolated and placed in ice-cold, Ca^2+^-free physiological saline solution (Mg-PSS), containing 140 mM NaCl, 5 mM KCl, 2 mM MgCl_2_, 10 mM HEPES, and 10 mM glucose (pH 7.4, adjusted with NaOH), supplemented with 0.5% bovine serum albumin (BSA). Cerebral pial arteries (anterior cerebral, middle cerebral, posterior cerebral, and superior cerebellar) were gently removed from the brain and washed in Mg-PSS. Third- to fifth-order mesenteric arteries (lumen diameter < 150 µm) were selected after removal of adipose, veins, lymph vessels, and connective tissue. Gracilis arteries were isolated from hind limbs and used for skeletal muscle resistance artery experiments.

Single native SMCs for patch-clamp electrophysiology and imaging experiments were obtained by initially digesting isolated arteries in 1 mg/mL papain (Worthington Biochemical Corporation), 1 mg/mL dithiothreitol (DTT), and 1 mg/mL BSA in Mg-PSS at 37°C for 10 min, followed by a 14-min incubation with 1 mg/mL type II collagenase (Worthington Biochemical Corporation). A single-cell suspension was prepared by washing digested arteries three times with Mg-PSS and triturating with a fire-polished glass pipette. All cells used for this study were freshly dissociated on the day of experimentation.

### Isolation of RNA and ddPCR

Total RNA was extracted from isolated tissue arteries by homogenization in TRIzol reagent (Invitrogen), followed by purification using a Direct-zol RNA microprep kit (Zymo Research) with on-column DNAse treatment. RNA concentration was determined using an RNA 6000 Pico Kit run on a Bioanalyzer 2100 using Agilent 2100 Expert Software (B.02.11; Agilent Technologies). RNA was converted to cDNA using iScript cDNA Supermix (Bio-Rad, Hercules). Quantitative ddPCR was performed using QX200 ddPCR EvaGreen Supermix (Bio-Rad), cDNA templates, and custom-designed primers. The primer sequences for *Trpm4*, *Trpv1*, *Trpc6*, *Ano1*, and *Pkd2* are provided in Table S1. Generated droplet emulsions were amplified using a C100 Touch Thermal Cycler (Bio-Rad), and the fluorescence intensities of individual droplets were measured using a QX200 Droplet Reader (Bio-Rad) running QuantaSoft (version 1.7.4; Bio-Rad). Analyses were performed using QuantaSoft Analysis Pro (version 1.0596; Bio-Rad).

### Fluorescence imaging

Intact tissues, vessels, and isolated native SMCs from *Trpm4-*Cre::mTmG reporter mice were imaged using a Nikon A1R HD laser scanning confocal microscope. The system is built onto an inverted Nikon Ti2-E Microscope. The illumination system comprises 488 nm and 561 nm diode laser lines, coupled to 2 GaAsP and 2 high-sensitivity Photomultiplier Tubes. Intact vessels were imaged with a 10 × /0.45 NA air objective (pixel size 0.8512 µm × 0.8512 µm; Plan Apochromat Lambda, Nikon). Isolated SMCs were imaged with a 60 ×/1.49 NA oil-immersion objective (pixel size 0.0496 µm × 0.0496 µm; Apochromat TIRF, Nikon). The microscope was controlled using the NIS-Elements C with JOBS Acquisition Module software. Representative images are shown as maximum-intensity projections of confocal Z-stacks unless otherwise indicated. For imaging of intact cerebral arteries in *Trpm4-*Cre::mTmG reporter mice, brains were rapidly removed and placed in a 35-mm glass-bottom dish containing PSS. Images were obtained from cortical regions adjacent to the middle cerebral pial artery. Mesenteric arteries (3^rd^-5^th^ order) were carefully dissected, cleaned of surrounding adipose and connective tissue, and transferred to a glass-bottom dish containing PSS. Skeletal muscle arteries were isolated from the gracilis muscle using a similar procedure and prepared for imaging under identical conditions. Individual SMCs were isolated as described above and seeded for 20 min on a 35-mm glass-bottom dish containing Ca^2+^-PSS.

### Electrophysiological recordings

Native SMCs were enzymatically isolated, transferred to a recording chamber (Warner Instruments, Holliston, MA, USA), and allowed to adhere to glass coverslips for 20 min at room temperature. Currents were recorded using an Axopatch 200B amplifier (Molecular Devices, Sunnyvale, CA, USA) equipped with an Axon CV-203BU headstage and a Digidata 1440A digitizer. Data were acquired and analyzed with Clampex and Clampfit software (version 10.2, Molecular Devices). Recording electrodes (3–5 MΩ) were pulled using a P-97 micropipette puller (Sutter Instruments Laboratories, Long Island, NY, USA) and filled with appropriate pipette solutions, as detailed below.

Whole-cell TRPM4 currents were recorded in a bath solution containing 156 mM NaCl, 1.5 mM CaCl_2_, 10 mM glucose, 10 mM Hepes, and 10 mM TEA-Cl (pH 7.4, adjusted with NaOH). The pipette solution contained 156 mM CsCl, 8 mM NaCl, 1 mM MgCl_2_, and 10 mM Hepes (pH 7.2, adjusted with CsOH), with free [Ca^2+^] buffered to 200 µM using CaCl_2_ and EGTA, as calculated with MaxChelator (WEBMAXC standard). Whole-cell cation currents were elicited by 400-ms voltage ramps (–100 to +100 mV) from a holding potential of –60 mV, repeated every 2 s for 300 s. The selective TRPM4 inhibitor NBA (10 µM) was applied after the peak current was reached (∼100 s). Current amplitude was expressed as the current at +100 mV. Summary current–voltage (I–V) plots were generated using values obtained from the last 50 ms of each step.

Stretch-induced currents were recorded as TICCs using the amphotericin B perforated-patch configuration. The bath contained 134 mM NaCl, 6 mM KCl, 1 mM MgCl_2_, 2 mM CaCl_2_, 10 mM Hepes, and 10 mM glucose (pH 7.4, adjusted with NaOH). The pipette solution contained 110 mM K-aspartate, 1 mM MgCl_2_, 30 mM KCl, 10 mM NaCl, 10 mM Hepes, and 5 µM EGTA (pH 7.2, adjusted with KOH), supplemented with amphotericin B (200 µg/mL) to enable electrical access. TICCs were recorded at a holding potential of –70 mV. Membrane stretch was produced by applying negative pressure (typically −20 mmHg) through the recording pipette using a pressure-clamp system (HSPC-1; ALA Scientific Instruments, Farmingdale, NY, USA). TICC activity was quantified as the sum of the open-channel probability (NP_o_) of multiple 1.75-pA open states^15^.

### Pressure myography

Pressure myography experiments were conducted in accordance with current guidelines^27^. Arteries were mounted between two glass cannulas (outer diameter, ∼40–50 µm) in a pressure myography chamber (Living Systems Instrumentation) and secured with a nylon thread. Intraluminal pressure was controlled using a servo-controlled peristaltic pump (Living Systems Instrumentation). Preparations were visualized with an inverted microscope (Accu-Scope Inc.) coupled to a USB camera (The Imaging Source LLC). Changes in luminal diameter were continuously recorded using IonWizard software (version 7.2.7.138; IonOptix LLC, Westwood, MA, USA). Cerebral arteries were bathed in warmed (37°C), oxygenated (21% O_2_, 6% CO_2_, 73% N_2_) artificial cerebrospinal fluid (aCSF; 124 mM NaCl, 3 mM KCl, 2 mM MgCl_2_, 2 mM CaCl_2_, 1.25 mM NaH_2_PO_4_, 26 mM NaHCO_3_, and 4 mM glucose) at an intraluminal pressure of 5 mmHg. For mesenteric arteries and skeletal muscle arteries, the chamber and cannula were filled with vessel PSS (119 mM NaCl, 4.7 mM KCl, 1.17 mM MgSO_4_, 1.8 mM CaCl_2_, 1.18 mM KH_2_PO_4_, 5 mM glucose, and 0.03 mM EDTA). Following equilibration for 15 min, intraluminal pressure was increased to 110 mmHg, and vessels were stretched to their approximate *in vivo* length, after which pressure was reduced back to 5 mmHg for an additional 15 min. The viability of each preparation was assessed by evaluating vasoconstrictor responses to high extracellular [K^+^] PSS, made isotonic by adjusting [NaCl] and [KCl] (67 mM NaCl/60 mM KCl for cerebral arteries, 63.7 mM NaCl/60 mM KCl for mesenteric and skeletal muscle arteries). Arteries that showed less than 10% constriction in response to elevated [K^+^] were excluded from further investigation.

Vessels were pressurized by increasing intraluminal pressure in 20 mmHg increments from 5 mmHg to physiologically relevant pressures (≥ 40 mmHg, depending on vessel response), and the active diameter was recorded after allowing vessels to equilibrate for at least 5 min or until a steady-state diameter was reached. Acute effects of NBA (10 µM) on myogenic tone were assessed by applying NBA while maintaining constant intraluminal pressure, and reversibility was determined by washout experiment. After washout of the NBA effect, arteries were superfused with Ca^2+^-free aCSF or Ca^2+^-free PSS supplemented with EGTA (2 mM) and the voltage-dependent [Ca^2+^] channel blocker diltiazem (10 µM) to inhibit SMC contraction, after which passive diameter was obtained. Myogenic tone was calculated as myogenic tone (%) = (1 – active or drug-treated lumen diameter/passive lumen diameter) × 100.

Noncumulative concentration-response curves were generated by applying increasing concentrations of NBA (10 nM, 100 nM, 300 nM, 1 µM, 3 µM, 10 µM, 30 µM) under steady-state pressurized conditions. Myogenic tone at each concentration was normalized to baseline tone prior to drug application.

To assess the full pressure-induced myogenic response, intraluminal pressure was increased in 20 mmHg increments from 5 mmHg to 140 mmHg. The active diameter was obtained by allowing vessels to equilibrate for at least 5 min at each pressure as mentioned above. The contribution of TRPM4 to pressure-induced tone was determined by first subjecting arteries to the pressure-step protocol under control conditions. Arteries were then returned to 5 mmHg and bathed with the selective TRPM4 inhibitor NBA (10 µM) before repeating the pressure-step protocol. Following completion of the pressure–response study, intraluminal pressure was lowered to 5 mmHg, and arteries were also superfused with Ca^2+^-free aCSF or Ca^2+^-free PSS supplemented with EGTA (2 mM) and the voltage-dependent [Ca^2+^] channel blocker diltiazem (10 µM), after which passive diameter was obtained by repeating the stepwise increase in intraluminal pressure.

### Statistical analysis

All summary data are presented as means ± SEM. GraphPad Prism software (Version 10.4.1) was used to conduct statistical analyses and generate graphs. The value of n refers to the number of cells for patch-clamp electrophysiological experiments or the number of arteries used for pressure myography experiments. Statistical analyses were performed using paired *t*-test, unpaired *t*-test, one-way analysis of variance (ANOVA) or two-way ANOVA, as appropriate. IC_50_ values for NBA were calculated using a non-linear regression analysis of concentration response curves. A *P*-value < 0.05 was considered statistically significant for all analyses.

## Author contributions

Conceptualization: SE, WZ

Methodology: SE, WZ, BL, SP, YFE

Investigation: WZ, BL, ASS, SP

Visualization: WZ, BL, SE

Funding acquisition: SE

Project administration: SE, YFE

Supervision: SE

Writing – original draft: SE, WZ

Writing – review & editing: SE, WZ, YFE

## Acknowledgments

This study was supported by R35HL155008 (to S.E.), R01DK135621 and R01HL122770 (to Y.F.E.).

**Figure Supplement 1. The selective AT_1_R antagonist losartan suppresses stretch-induced TICC activity but does not inhibit Ca^2+^-activated whole-cell TRPM4 current in arterial SMCs. A,** Conventional whole-cell patch-clamp recordings from cerebral artery SMCs. Left, representative current–voltage (I–V) relationship before and after application of losartan (10 µM). Right, summary current density measured at +100 mV. Data are means ± SEM (n = 6 cells from 4 mice; *P* = 0.1579; paired *t*-test). Symbols represent individual cells. **B**, Perforated-patch recordings from cerebral artery SMCs. Left, representative recording showing TICC activity at 0 mmHg, during membrane stretch induced by −20 mmHg suction, and after application of losartan (10 µM). Right, summary TICC activity expressed as NP_o_. Data are means ± SEM (n = 6 cells from 5 mice; *P* < 0.0001 for 0 mmHg vs. −20 mmHg, *P* < 0.0001 for −20 mmHg vs. losartan, and *P* = 0.8605 for 0 mmHg vs. losartan; repeated-measures one-way ANOVA with Tukey test). Symbols represent individual cells. **C**, Conventional whole-cell patch-clamp recordings from mesenteric artery SMCs. Left, representative I–V relationship before and after losartan. Right, summary current density measured at +100 mV. Data are means ± SEM (n = 5 cells from 5 mice; *P* = 0.5053; paired *t*-test). Symbols represent individual cells. **D**, Perforated-patch recordings from mesenteric artery SMCs. Left, representative recording showing TICC activity at 0 mmHg, during −20 mmHg suction, and after losartan. Right, summary TICC activity expressed as NP_o_. Data are means ± SEM (n = 7 cells from 4 mice; *P* < 0.0001 for 0 mmHg vs. −20 mmHg, *P* < 0.0001 for −20 mmHg vs. losartan, and *P* = 0.9905 for 0 mmHg vs. losartan; repeated-measures one-way ANOVA with Tukey test). Symbols represent individual cells. **E**, Conventional whole-cell patch-clamp recordings from skeletal muscle artery SMCs. Left, representative I–V relationship before and after losartan. Right, summary current density measured at +100 mV. Data are means ± SEM (n = 6 cells from 2 mice; *P* = 0.1510; paired *t*-test). Symbols represent individual cells. **F**, Perforated-patch recordings from skeletal muscle artery SMCs. Left, representative recording showing TICC activity at 0 mmHg, during −20 mmHg suction, and after losartan. Right, summary TICC activity expressed as NP_o_. Data are means ± SEM (n = 5 cells from 3 mice; *P* = 0.0002 for 0 mmHg vs. −20 mmHg, *P* = 0.0010 for −20 mmHg vs. losartan, and *P* = 0.9902 for 0 mmHg vs. losartan; repeated-measures one-way ANOVA with Tukey test). Symbols represent individual cells.

## Notes

### Competing Interest Statement

The authors have declared no competing interest.

